# Blood and Neuronal Extracellular Vesicle Mitochondrial Disruptions in Schizophrenia

**DOI:** 10.1101/2025.07.30.667452

**Authors:** A. Ankeeta, Ashutosh Tripathi, Bindu Pillai, Yizhou Ma, Joshua J. Chiappelli, Jessica N. Jernberg, Keiko Kunitoki, Xiaoming Du, Si Gao, Bhim M. Adhikari, Consuelo Walss-Bass, Giselli Scaini, Peter Kochunov, Anilkumar Pillai, L Elliot Hong

## Abstract

The high energy demand of the human brain obligates robust mitochondrial energy metabolism, while mitochondrial dysfunctions have been linked to neuropsychiatric disorders including schizophrenia spectrum disorders (SSD). However, *in vivo* assessments that can directly inform brain mitochondrial functioning and its etiopathophysiological path to SSD remain difficult to obtain. We hypothesized that system and brain mitochondrial dysfunctions in SSD may be indexed by elevated cell-free mitochondrial DNA (cf-mtDNA) levels in the blood and in neuronal extracellular vesicles (nEVs). We also explored if these mtDNA marker elevations were associated brain metabolites as measured by magnetic resonance spectroscopy (MRS). We examined blood cf-mtDNA in 58 SSD patients and 33 healthy controls, followed by assessing nEV mtDNA and metabolite levels using MRS in a subgroup of patients and controls. We found that people with SSD had significantly elevated cf-mtDNA levels in both the blood (p=0.0002) and neuronal EVs (p=0.003) compared to controls. These mtDNA abnormalities can be linked back to brain lactate+ levels such that higher blood and nEV mtDNA levels were significantly associated with higher lactate+ levels measured at the anterior cingulate cortex (r=0.53, 0.53; p=0.008, 0.03, respectively) in SSD patients. Furthermore, higher developmental stress and trauma were significantly associated with higher cf-mtDNA levels in both the blood and neuronal EVs in SSD patients (r=0.29, 0.49; p=0.01, 0.03, respectively). In conclusion, if replicated and fully developed, blood and neuronal EV-based cell free mtDNA may provide a clinically accessible biomarker to more directly evaluate the mitochondrial hypothesis and the abnormal bioenergetics pathways in schizophrenia.

## Introduction

The etiopathophysiology of schizophrenia spectrum disorders (SSD) remains elusive. The mitochondrial hypothesis on SSD has emerged and been gaining evidence [1–6]. Mitochondrion is maternally inherited with circular double-strand mitochondrial DNA (mtDNA) and the primary organelle for energy production, among serving other cellular functions [7]. The human brain has a high rate of metabolism, making mitochondrion a focus for the research on brain health and diseases [6]. While not consistent, evidence directly involving the mitochondrion in SSD patients includes mitochondrial gene expression, mtDNA mutation or copy number abnormalities in postmortem brain tissues [8,9], reprogrammed or enriched patient neuronal cells [2], and peripheral blood samples [10,11]. Other direct evidence includes mitochondrial electron transport chain dysfunction [3,12], and mitochondrial morphological alternations or reduced mitochondrial numbers [6,13]. There is also strong indirect evidence inferring mitochondrial involvement in SSD by demonstrating energy metabolism abnormalities in the brain shown by ^31^P-MRS [4,5]. However, none of these methods directly assess neural mitochondrial abnormalities in clinical patients, hindering the progress in fully testing the mitochondrial involvement in SSD.

Each cell contains hundreds to thousands of mitochondria, which are highly dynamic and frequently move, fuse, divide, and breakdown for rapid adaptation and damage repair in order to meet energetic demands [13]. A byproduct of these dynamic mitochondrial activities is cell-free mtDNA (cf-mtDNA), measurable in the circulation either as passive leakages or active removal of dysfunctional mitochondria for quality control in basal but particularly in stressful conditions [14]. Cell-free but functional mtDNA may also serve intercellular exchange functions although this claim remains controversial [15,16]. What is well-established is that mitochondrial dysfunctions increase the release of mitochondrial contents including mtDNA [17]. Studies have shown that patients with SSD have elevated plasma levels of cf-mtDNA [18,19], which progressively decreases following antipsychotic treatment [20]. Therefore, abnormal cf-mtDNA levels support the mitochondrial dysfunction hypothesis for SSD. However, blood cf-mtDNA can be originated from any organ. Accordingly, our study also assessed whether the abnormal cf-mtDNA levels have a neural origin using neuronal extracellular vesicles (nEVs).

EVs are small-sized exosomes to large-sized micro-vesicles that are produced by cells from all organs to transport nanoparticle cellular contents needed for critical functions including the removal of damaged organelles and intercellular communications [21]. A small portion of EVs in the blood are produced by neural cells in the brain, providing a rare opportunity for *in vivo* bio-sampling of brain contents. nEVs have been increasingly applied to SSD research [22]. Neurons can exude EVs that contain functional or damaged mitochondria or their contents [23]. Initial EV-based mitochondrial studies in SSD showed that several key mitochondrial proteins are either elevated or reduced and mitochondrial ATP production is reduced in a modest sample of 10 first episode patients [24]. We are unaware of any previous studies of cf-mtDNA from EVs of neuronal origin in SSD.

To further test that the cf-mtDNA measured from nEVs are related to brain functioning, we explored the relationship between cf-mtDNA and brain metabolites as measured by ^1^H magnetic resonance spectroscopy (MRS), as several metabolites measurable by ^1^H MRS such as lactate and N-acetylaspartate (NAA) are important bioenergentics involved in mitochondrial functions [25,26], and we have recently showed that greater stressor exposure was associated with brain lactate and NAA in SSD [27]. Studies have observed elevated lactate levels in the brains of patients with SSD although the results are inconsistent [28–33]. To our knowledge, no study has attempted to link brain-based metabolites and neuronal EV-based mitochondrial functions in SSD-related conditions. The anterior cingulate cortex (ACC) was selected for MRS acquisition in part due to its central role in emotion regulation, cognition, and its established vulnerability in SSD. Specifically, decreased mitochondrial density in the postmortem ACC of patients with schizophrenia has been reported [34], and astrocyte-derived lactate in the ACC regulates neuronal mitochondrial biogenesis [35], providing additional justification to explore the relationship between cf-mtDNA and brain metabolites in the ACC region.

Finally, chronic stress can alter energy metabolism, causing mitochondrial structural and functional disruption and mtDNA release [36,37]. Stress from maternal exposure to adverse life events in pregnancy [38,39] to early childhood adversities [40] is known to increase the risk for SSD [41,42]. Using animal models, we had shown that chronic stress can drastically increase cf-mtDNA release, suggesting that cf-mtDNA may provide a biomarker to index the stress-mitochondrial pathway dysfunction [43]. Whether this stress – cf-mtDNA relationship is present in humans or abnormal in SSD patients is unknown. As stress exposure during early development can impact mitochondrial enzyme functioning later in adulthood in animal models [44–46], this study assessed whether developmental and/or recent stressors may contribute to cf-mtDNA in SSD.

## Materials and Methods

### Participants

The study included 91 participants who completed blood mtDNA assays: N=58 people with schizophrenia spectrum disorders (SSD) (including 49 schizophrenia, 8 schizoaffective disorder, and 1 schizophreniform disorder) and N=33 healthy controls (**Table 1**). Patients were recruited from the outpatient clinics of the Maryland Psychiatric Research Center and several nearby mental health clinics. Healthy controls (HC) were recruited through local media advertisements. Structured Clinical Interview for DSM-4 or 5 was used to confirm or exclude psychiatric diagnoses. Exclusion criteria included a history of neurological conditions, head trauma with cognitive sequelae, and active and uncontrolled major medical conditions. Participants with substance abuse or dependence other than nicotine or cannabis in the 6 months prior to study were excluded. Chlorpromazine dose equivalent (CPZ) was calculated for each patient’s antipsychotic medication regimen at the time of the study [47]. All patients were on antipsychotic medications, including 7 taking typical antipsychotics, 47 taking atypical (including 34 taking clozapine), and 4 taking a combination of typical and atypical antipsychotics. The mean CPZ was 526.8 mg, with a range of 100–3066 mg daily. Data were collected under protocols approved by the University of Maryland Institutional Review Board. Participants provided written informed consent before participation.

**Table 1.**
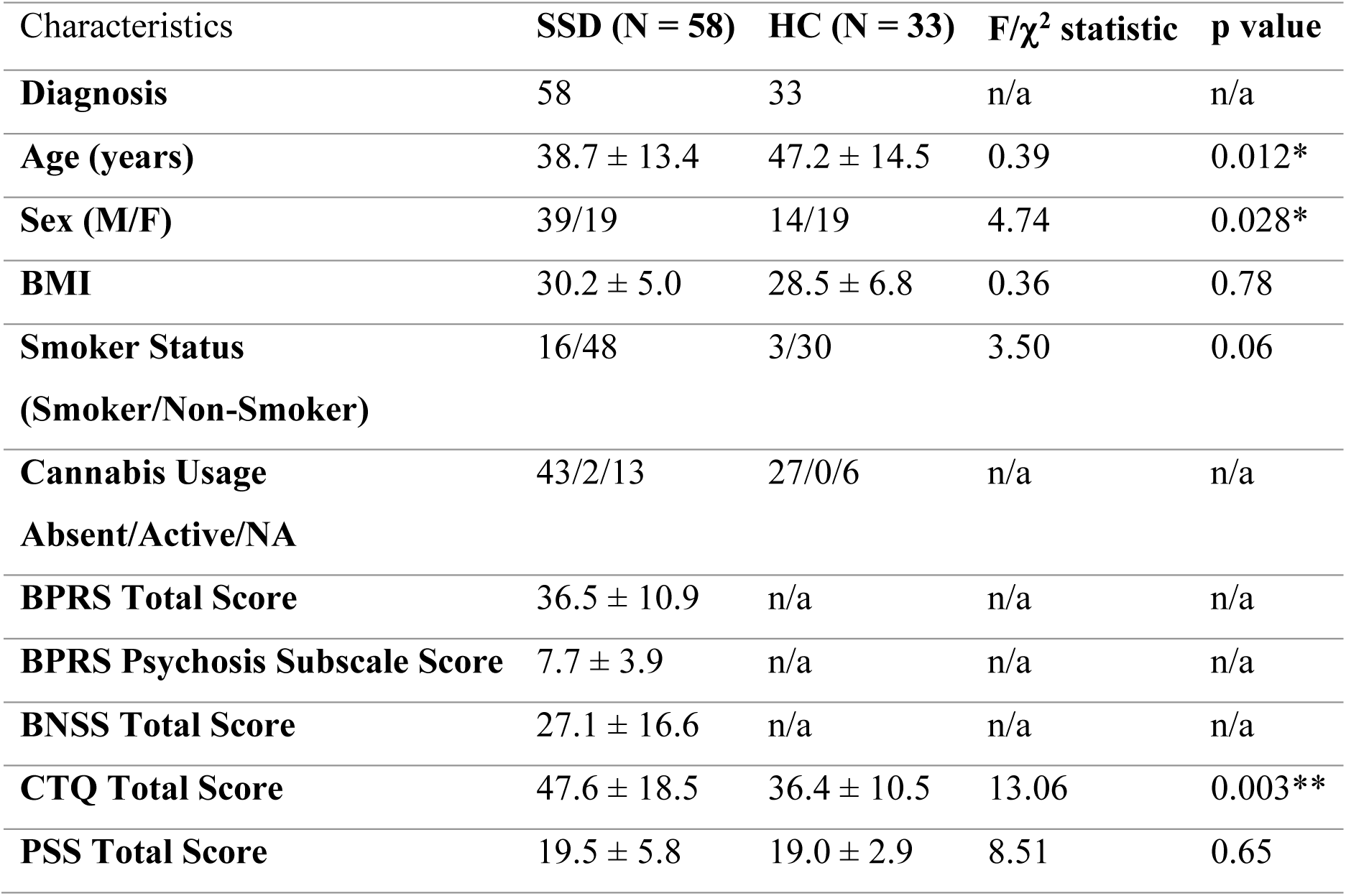
Demographic and clinical characteristics between patients with schizophrenia spectrum disorder (SSD) and healthy controls (HC). BPRS: Brief psychiatric rating scale. BNSS: Brief negative symptom scale. CTQ: Childhood trauma questionnaire. PSS: Perceived stress scale. *p<0.05, **p<0.001. NA: not available or not applicable.

### Clinical measures

Overall psychiatric symptoms and negative symptoms were assessed using the Brief Psychiatric Rating Scale (BPRS) [48] and Brief Negative Symptom Scale (BNSS) [49], respectively. Psychotic symptom severity was measured by the BPRS psychosis subscale. The Childhood Trauma Questionnaire - Short Form (CTQ) was used to assess developmental stressful experiences consisting of physical, emotional, and sexual abuse, and neglect during childhood [50]. The Perceived Stress Scale (PSS) was used to measure subjective levels of stress over the past month [51].

### Magnetic Resonance Spectroscopy

Proton magnetic resonance spectroscopy (¹H-MRS) was conducted using a single-voxel short-echo PRESS sequence on a Siemens MAGNETOM 3T scanner to assess neurometabolite concentrations. The anterior cingulate cortex (ACC) voxel measuring 40×30×20mm³ (24cm³) and was manually positioned on the midline in the dorsal ACC, oriented to minimize volume effects and lipid contamination. The sequence parameters were optimized for short-TE acquisition (TE=30ms) that provides high signal-to-noise ratio and minimal T2 signal decay, TR=2000 ms, flip angle=90°, 128 signal averages, 2048 data points, and bandwidth=2000 Hz, with an acquisition time of 4 minutes and 24 seconds. Water suppression was enabled using frequency-selective saturation pulses (bandwidth=50 Hz), and unsuppressed water reference spectra were also collected for quantification and eddy current correction. The sequence incorporates automatic phase cycling, gradient spoiling, and oversampling removal to enhance spectral fidelity and reduce contamination from outer voxel regions. The short TE (30 ms) does not separate metabolites like lactate from lipid and other contaminants as compared to longer TEs (e.g., 144 ms) or editing sequences (e.g., MEGA-PRESS) that are typically needed to increase lactate specificity from lipid and threonine [52,53]. Meanwhile, the current sequence retains high signal-to-noise ratio across a wider metabolic range, and the putative lactate identified at 1.33 ppm by a characteristic doublet was quantified as lactate+ (lac+). Similar approaches using TE around 30 ms have successfully quantified lactate+ in clinical populations, including schizophrenia and bipolar disorder [54–59]. LCModel (http://s-provencher.com/lcmodel.shtml) was used for quantification using a simulated basis set of 19 metabolites. Metabolite levels were referenced to water and were reported in institutional units. Spectra with FWHM>0.1 ppm and signal to noise ratio<10 reported by LCModel were excluded from analyses. We used Cramér-Rao lower bounds (CRLB) %SD ≤ 20% for all metabolites except 30% for lactate+, as the 20% threshold may be too conservative for low concentration metabolites like lactate+ [27,29,32]. Parameters exceeding these criteria were considered outliers. The proportion of the cerebrospinal fluid (CSF) and white matter within the spectroscopic voxels was calculated based on segmentation of MP-RAGE images. Metabolite values were corrected by dividing raw values by (1-CSF fraction) except lactate+ [60,61] because CSF contains lactate and has been suggested as a biomarker for mitochondrial dysfunction in SSD [62]. The default macromolecule and lipid components basis set in the LCModel was used [63]. These macromolecule components are simulated and incorporated into the fitting model to account for baseline contributions [64]. Structural T1-weighted images for co-registration were acquired using a fast spoiled gradient recalled sequence (TR=11.08ms, TE=4.3ms, flip angle=45°, FOV=256mm, 256×256 matrix, 172 slices, 1mm³ resolution) to obtain masks for ACC voxel placement. Twenty-two patients with SSD and 17 HC had MRS completed; after 2 patients and 1 control was excluded as outliers, 20 SSD and 16 HC were included for the analysis. An example of voxel placement and spectra is shown in **Figure 2A**.

### Blood plasma/serum mtDNA analysis by qRT-PCR

The sample included 58 SSD patients and 33 controls with blood samples available. Blood samples were collected in the morning hours following an overnight fast, which is relevant as circulating cf-mtDNA levels may exhibit diurnal fluctuations [65]. Among them, 30 had only serum, 32 had only plasma, and 29 had both serum and plasma available. For plasma samples, whole blood was collected in EDTA-containing tubes (Vacutainer), which were immediately centrifuged at 1000×g, 10 min. For serum samples, additional whole blood was collected using anticoagulant-free vacuum tube and centrifuged at 3800 rpm for 10 min. Samples were then aliquoted into separate tubes and stored at −80 °C until assay. Total cell-free DNA was isolated from 200μl plasma and/or serum using DNA isolation/purification kits (Qiagen, Hilden, Germany) according to the manufacturer’s protocol. Quantity of cf-mtDNA was measured by qRT-PCR by analyzing mitochondrially encoded NADH dehydrogenase subunit 1 (ND1) (forward primer-5’ ATACCCATGGCCAACCTCCT 3’ and reverse primer-5’ GGGCCTTTGCGTAGTTGTAT 3’), and NADH dehydrogenase subunit 4 (ND4) (forward primer-5’ TCTTCTTCGAAACCACACTT 3’ and reverse primer-5’ AAGTACTATTGACCCAGCGA 3’) genes. Further details are in Supplementary Information (**SI1**). Analysis of the mtDNA was done using the 2 (-delta delta Ct) method [66,67]. Cycle threshold (Ct) values were collected for all targets in each sample after the qRT-PCR run. Delta Ct values were calculated by subtracting the Ct values of mtDNA from the nDNA. Delta delta Ct was then calculated by subtracting delta Ct of the SSD patients from the delta Ct of the controls. Relative quantity (RQ) of mtDNA was calculated using 2^delta delta Ct [43,68,69]. MtDNA from neuronal EV used the same methods once neuronal EVs were isolated (see below).

### Isolation and validation of neuronal EVs

EVs were isolated from 0.5ml of human serum following a detailed protocol [77], details in **SI2**. To enrich for neuronal EVs, the suspensions were incubated at 4°C for 1 hour with 4 μg of biotinylated anti-human L1 cell adhesion molecule (L1CAM) antibody as the neuronal pulldown method following standard protocol (see **SI3** for detail). To analyze mtDNA levels in neuronal EVs, we extracted total DNA from neuronal EVs using the DNeasy Blood and Tissue Kit (Qiagen, Hilden, Germany). Quantification of mtDNA was performed using qRT-PCR, following the same methodology as used for blood plasma/serum mtDNA analysis. We then performed validations on the efficiency of the L1CAM pulldown using several steps, details in **S13**. These experiments confirmed the validity and efficiency of the L1CAM pulldown of the neuronal EV (see Results).

### Statistics

Group differences on key variables were tested using analysis of covariance (ANCOVA) with age and sex as covariates. Pearson’s correlations or linear regression analyses were used to determine associations between cf-mtDNA and stress and MRS measures, with a specific focus on those measures where the significant associations can be replicated in blood and neuronal cf-mtDNA in the expected direction. All tests were two-tailed. All statistical analyses were conducted using IBM SPSS Statistics version 28. Group differences in cf-mtDNA (blood and neuronal EVs) were tested using analysis of covariance (ANCOVA), with age and sex included as covariates. For within-group associations between cf-mtDNA levels and metabolite concentrations (e.g., lactate+) or clinical variables (e.g., childhood trauma), we used Pearson’s correlation or multiple linear regression models adjusting for age and sex. All statistical tests were two-tailed with alpha=0.05.

## Results

### Increased blood cf-mtDNA levels in SSD patients

The relative quantity (RQ) for ND1 and ND4 were fully complementary (r=1.0) in this sample, and thus all analytic results are identical using either ND1 or ND4; therefore, all subsequent analysis used ND1 data only to represent mtDNA. We first examined participant data with both serum and plasma cf-mtDNA levels available, and found that there were no significant differences between serum and plasma RQ (p=0.19) for ND1. Therefore, we combined the serum and plasma data by using plasma data when available and if not, serum data were used, and described the results as blood cf-mtDNA. Blood cf-mtDNA levels were significantly elevated in SSD patients compared to HC without age and sex correction (F=15.04, p=0.0002). The SSD group was younger (39.4±13.4 years) compared to controls (47.2±14.5; p=0.012) and had a higher proportion of males (39M/19F) than the control group (14M/19F; p=0.028) (**Table 1**). Adjusting for age and sex, the group differences in blood cf-mtDNA levels were significant (F=15.07, p=0.0002) (**Figure 1A**).

**Figure 1.**
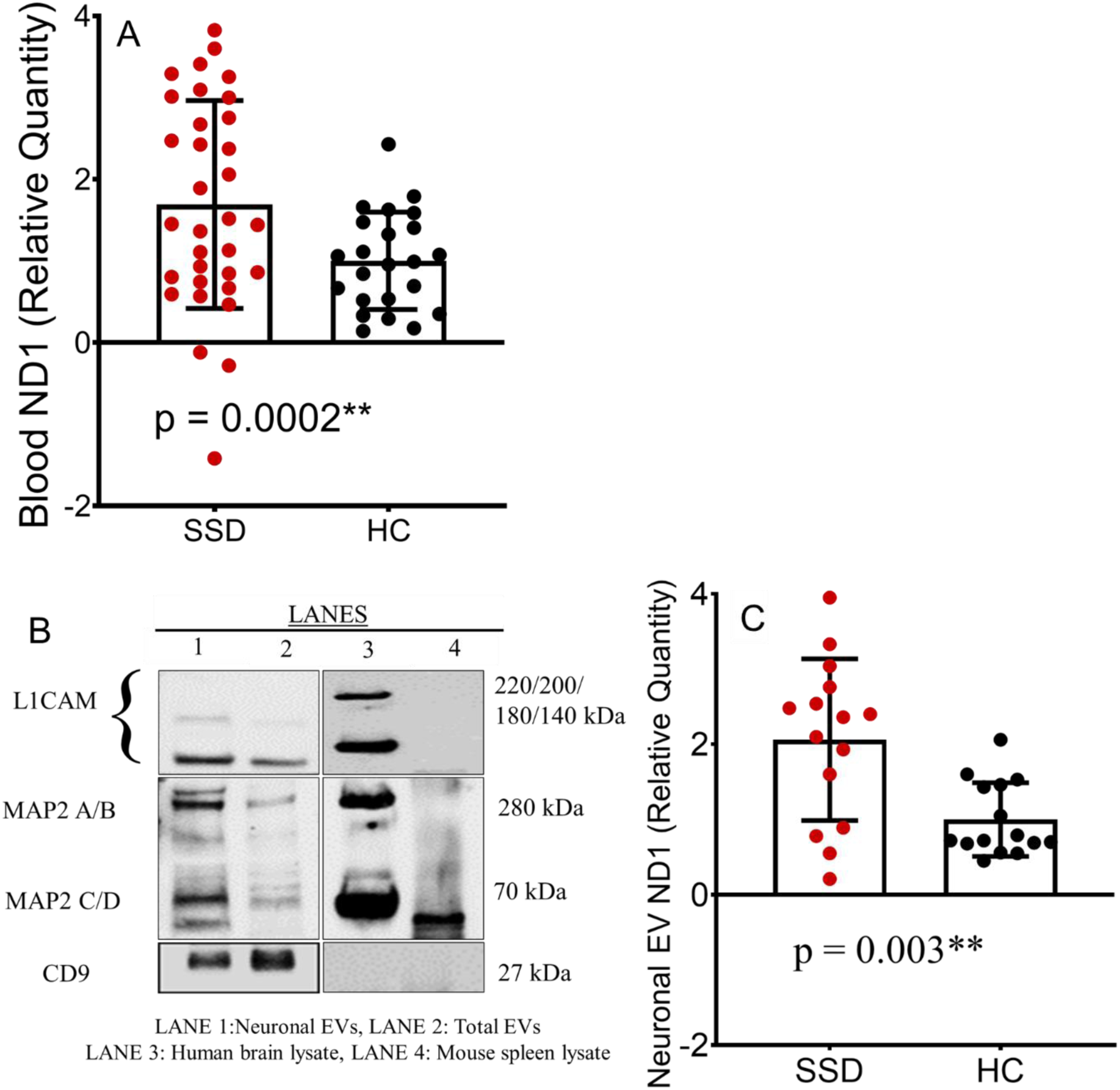
Cell free Blood mtDNA and neuronal EV characterization in SSD as compared to healthy controls. (A) Box plot comparing blood cf-mtDNA levels between patients with schizophrenia spectrum disorder (SSD, red) and healthy controls (HC, black). SSD patients exhibited significantly higher blood cf-mtDNA levels compared to HC, controlling for age and sex. Error bars indicate standard deviations (B) Western blot analysis demonstrating the enrichment of neuronal markers—neural cell adhesion molecule (L1CAM) and microtubule associated protein-2 (MAP-2)—in neuronal extracellular vesicles (EVs) (Lane 1) compared to total EVs (Lane 2) isolated from the serum of a single healthy control. Human brain lysate (Lane 3) and mouse spleen lysate (Lane 4) served as positive and negative controls, respectively, to confirm antibody specificity. CD9 was included as a canonical EV marker. (C) Box plot comparing neuronal EV cf-mtDNA levels between SSD and HC groups.

### Increased neuronal extracellular vesicle (nEV) cf-mtDNA levels in SSD patients

To support that the blood cf-mtDNA finding, we extracted nEVs from a subsample of 15 SSD (age 40.9±14.4, 10 males) and 15 HC (age 45.4±15.8, 7 males) from the above sample by frequency-matching their age (p=0.42) and sex (p=0.46). We first tested the neuronal nature of the EV, and found significantly higher levels of L1CAM in the neuronal EVs as compared to the total EVs, confirming the efficiency of the L1CAM pulldown method for neuronal EVs. Further, both isoforms of MAP2, a well-established neuronal marker, were detected at higher levels in the nEVs and human brain lysate compared to the total EVs (LANE 1, 2 and 3, **Figure 1B**), confirming neuronal enrichment of the nEVs; while mouse spleen lysate (LANE 4) lacked detectable expression of these markers. Overall, these results confirmed the primarily neuronal nature of the nEVs and validated the efficacy of L1CAM-mediated pulldown in isolating nEVs. We then measured cf-mtDNA from these nEVs, and found that nEV cf-mtDNA levels were significantly elevated in SSD patients compared to controls (F=11.1, p=0.003) (**Figure 1C**).

### Anterior cingulate metabolites levels and associations with cf-mtDNA levels

No significant differences in anterior cingulate cortex (ACC) metabolite levels were observed between SSD patients and HC after controlling for age and sex (**Figure 2B**). cf-mtDNA showed nominally significant correlations only with lactate+ (r = 0.53, p = 0.008) in the patients but not in controls; while nominally significant correlations with NAA in the controls (r = –0.44, p = 0.042) but not in patients (**Figure 2C**). However, there were no significant group differences in the strength of these correlations between SSD and HC.

**Figure 2.**
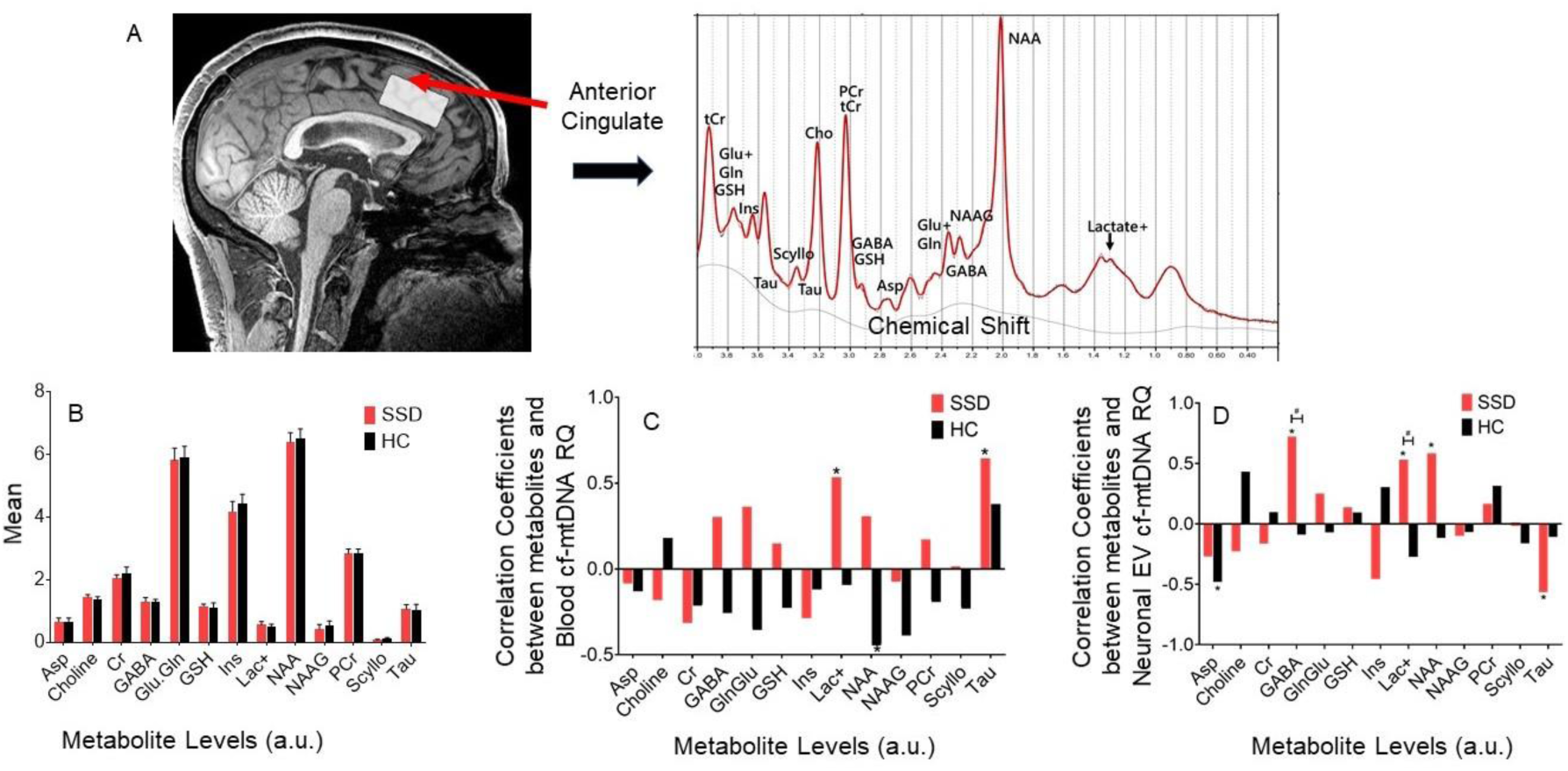
**A**: Voxel placement and representative MRS spectra from the anterior cingulate cortex (ACC), along with the corresponding LCModel-derived metabolite identification graph. **B**: Group comparisons of ACC metabolite levels between patients and healthy controls. Group differences were illustrated in **C** and **D** for correlation coefficients between ACC metabolites and cf-mtDNA levels in blood and neuronal extracellular vesicles (EVs), respectively. All correlations were controlled for age and sex. Error bars represent standard deviations.

In the subsample that had both spectroscopy and nEV data available (15 SSD and 14 HC), after controlling for age and sex, significant correlations were observed between nEV cf-mtDNA and GABA (r=0.72, p=0.003), NAA (r=0.58, p=0.018), and lactate+ (r=0.53, p=0.032) in SSD and aspartate in HC (r=–0.48, p=0.041) (**Figure 2D**). Overall, only lactate+ showed significant associations with cf-mtDNA that was replicated between blood (**Figure 3A**) and nEV cf-mtDNA in SSD (**Figure 3C**) but not in HC (**Figure 3B** and **3D**). To partially examine for specificity, we also extracted the lipid spectra that may have overlapped with the lactate+ spectra, and found that none of the lipid spectra in either patients or controls showed a significant association with blood or nEV cf-mtDNA (see **SI4**), supporting the likely lactate effects in the lactate+ associations with cf-mtDNA. Of note, the blood and nEV cf-mtDNA levels were positively correlated, similarly in patients (r=0.45, p=0.061), controls (r=0.56, p=0.017), and the combined sample (r=0.57, p=0.001).

**Figure 3.**
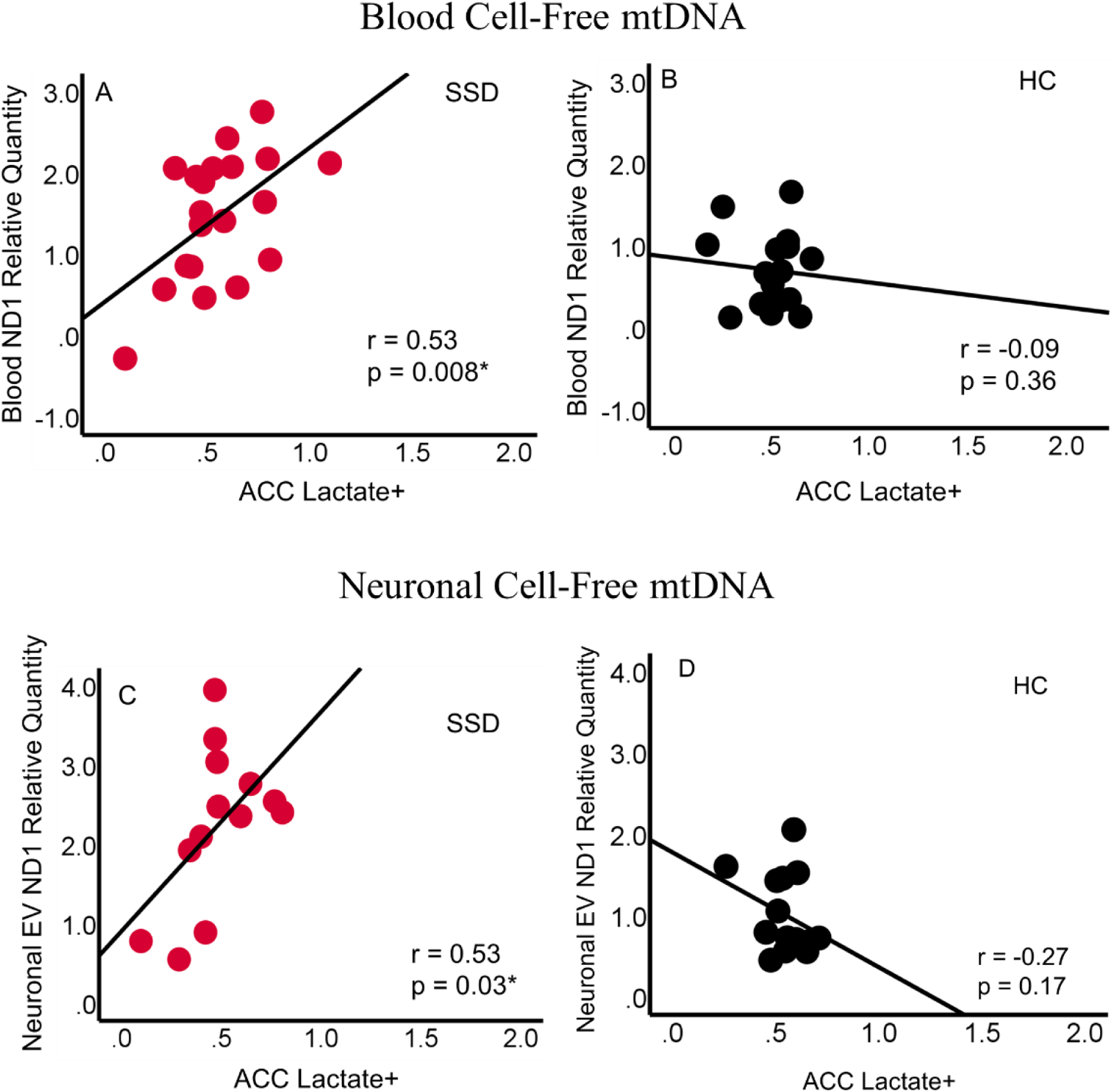
Associations between blood and neuronal EV cf-mtDNA levels and lactate concentrations: Blood cf-mtDNA was significantly associated with lactate+ levels in the anterior cingulate cortex in patients (A), but not in controls (B). Similarly, neuronal EV cf-mtDNA showed a significant correlation with lactate+ in patients with SSD (C), but not in controls (D).

### Stress and clinical associations with blood and nEV cf-mtDNA levels

Patients with SSD reported significantly higher CTQ total scores compared to HC (p=0.003). Higher CTQ total score was significantly associated with blood cf-mtDNA levels in patients (r=0.29, p=0.017) (**Figure 4A**), indicating that early life stress may contribute to elevated cf-mtDNA levels in adult patients. We found a similar but not significant trend in controls (r=0.27, p=0.069) (**Figure 4B**). Higher CTQ total score was also significantly associated with nEV cf-mtDNA (r=0.49, p=0.032) in SSD but not HC (r=−0.10, p=0.36) (**Figure 4C-D**). In comparison, The perceived stress scale total score (recent month) was not significantly associated with either blood (r=0.14, p=0.17) or nEV cf-mtDNA (r=−0.014, p=0.48) in the patients or the controls (r=0.12, p=0.26 and r=0.38, p=0.90, respectively).

**Figure 4.**
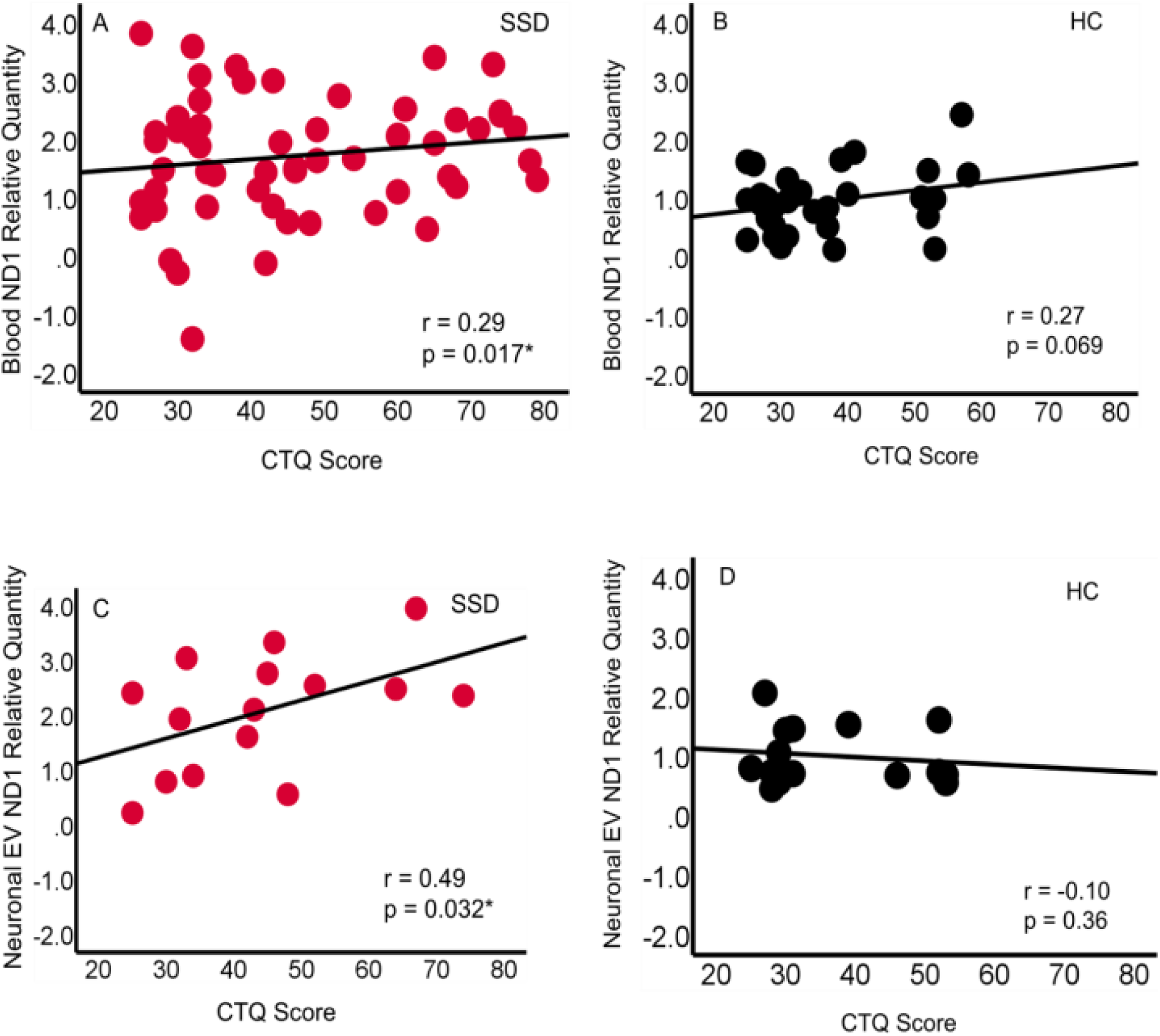
Higher childhood stress significantly contributed to elevated blood and neuronal extracellular vehicle cf-mtDNA in patients (**A, C**), as indicated by significant correlations. This effect was not significant in healthy controls (**B, D**).

Regarding relationships between cf-mtDNA and clinical symptoms and treatment in patients, the BPRS total score was not significantly associated with blood (r=−0.02, p=0.44) or nEV (r=−0.09, p=0.38) cf-mtDNA RQ. BPRS psychosis subscale was also not significantly associated with blood (r=−0.19, p=0.91) or nEV (r=−0.23, p=0.21) cf-mtDNA. Negative symptoms as measured by the BNSS was significantly and negatively associated with blood (r=−0.39, p=0.003) but not in nEV (r=−0.13, p=0.33) cf-mtDNA. We examined the relationship between perceived stress levels (measured by the Perceived Stress Scale, PSS) and mtDNA levels in both blood and nEVs. PSS scores showed no significant correlation with blood (r=0.14, p=0.17; r=0.13, p=0.27) or neuronal cf-mtDNA levels (r=−0.14, p=0.48; r=0.38, p=0.09) in the SSD and HC groups, respectively. BMI was not significantly different between groups (**Table 1**); there was no significant correlation between BMI and either blood or neuronal mtDNA in SSD or HC (all p>0.05); and adding BMI as a covariate did not meaningfully change the group difference statistics. Many antipsychotic medications can inhibit the mitochondrial respiratory chain and damage the mitochondria [71,72], but we found that CPZ was non-significantly associated with blood (r=−0.07, p=0.33) or nEV (r=−0.27, p=0.36) cf-mtDNA. Comparing cf-mtDNA levels in clozapine vs. non-clozapine patients, we found no significant differences in blood (p=0.69) or nEV (p=0.08) cf-mtDNA, where patients on clozapine showed numerically lower nEV cf-mtDNA levels. Finally, tobacco smoking contains high levels of reactive oxygen and nitrogen species thought to contributed to oxidative stress and mitochondria damages [73,74], but we found that current smokers had no significant differences in blood (F=0.12, p=0.73) and nEV cf-mtDNA (F=1.04, p=0.35) compared to non-smokers.

## Discussion

The study results showed that cell-free mtDNA levels in the blood and in neuronal EVs are significantly elevated in patients with SSD compared to healthy controls, and these mitochondrial indices measured in the periphery were significantly associated with lactate+ levels measured at the anterior cingulate cortex in the patients. These findings provide replications of the blood cf-mtDNA abnormalities observed in previous studies that indicated mitochondrial dysfunctions and more generally the energy metabolism dysfunctions in SSD. Two primary novel findings of the study are that the abnormal cf-mtDNA levels can be identified in nEV, and that this cf-mtDNA abnormality in SSD is nominally linked to several brain metabolite levels, but replicable between blood and nEV cf-mtDNA only with the lactate+ measure. Furthermore, developmental traumatic experiences in childhood appeared to significantly contribute to elevated cf-mtDNA levels in adult SSD patients.

Increased blood cf-mtDNA has been reported in SSD [18,19], which may be attributed to more mitochondrial breakdown and released mtDNA or possibly also increased intercellular exchanges [17]. Our cf-mtDNA findings in the blood replicated these findings, although they may not reflect mitochondrial dysfunctions in the brain. To the best of our knowledge, cf-mtDNA in nEVs has never been reported in SSD or other psychiatric conditions. While L1CAM was initially thought as be as one of the best pulldown tagging antibodies for brain nEVs, some studies cast doubt on the validity [75]. More recent studies using carefully designed single EV assays did confirm that L1CAM-tagged EVs are primarily neuron-derived and of brain origin [76]. However, given the ongoing debates, we performed additional validation that supported the neuronal nature of the L1CAM-tagged EVs in this sample. The findings showed that neuronal cf-mtDNA release can be clinically measured and is likely elevated in patients with SSD, providing a potential novel biomarker to study the mitochondrial involvements in SSD.

The cause of the elevated cf-mtDNA in SSD is not entirely clear. In cancer, cardiovascular disease, aging, and other conditions, heightened mtDNA mutations may increase oxidative stress, mitochondrial fragmentations, mitophagy, and fragmented mtDNA release [77–81]. Increased mtDNA mutations have been reported in some initial SSD studies although the findings are not consistent, and many subsequent studies failed to find SSD to be associated with more mtDNA mutations [82–84]. Therefore, whether mtDNA mutations have contributed to increased cf-mtDNA in SSD requires additional investigations.

The association with brain lactate+ levels is a novel finding and may offer another potential mechanism for the elevated cf-mtDNA. The MRS sequence used in this study was to broadly survey multiple metabolites and the lactate+ spectra are likely contaminated with other components, thus the following discussion on lactate-related mechanisms should be viewed as speculative, but the significant correlation with lactate+ in patients may be consistent with the notion that impaired mitochondrial function can lead to elevated lactate [85–87]. Mitochondrial lactate dehydrogenase (LDH) facilitates tissue lactate clearance and oxidation in vivo [88], and excessive mitochondrial breakdown may reduce the clearance leading to higher lactate. This or other mechanisms may induce changes in lactate and mtDNA levels in the same direction and thus the significant association. It is possible that such association would not be apparent in normal conditions and can only be observed in a pathophysiologically perturbed mitochondrial condition as hypothesized in SSD, which could explain the significant association only in patients but not in healthy controls. However, the finding was based on a modest sample and replication is required, and additional studies are needed to support the speculated mechanisms.

Another likely source of the elevated cf-mtDNA in SSD is stress. Stressor exposure in early development can increase the risk for SSD [38–40,89]. Stress and other toxic processes can increase mitochondrial damages, and mechanisms to remove the superfluous mitochondria are important for mitochondrial quality control in stressful conditions [14]. Chronic stress in particular can exacerbate mitochondrial energy demand, disruption, and mtDNA release [36,37], especially the increase of cf-mtDNA release that has been associated with social behaviors and inflammatory processes [43]. Previously it was unclear if this mechanism may be applicable to SSD. Here, we observed that more childhood trauma is associated with elevated blood and neuronal cf-mtDNA in adult SSD patients, providing supportive evidence between past chronic stressors and current mitochondrial dysfunction in SSD. This finding also corroborates animal models where stressors applied early during development can deregulate mitochondrial enzyme functioning that persists into adulthood animals [44–46].

This study has several limitations. Patients are on antipsychotic medications, which have a complex effect on mitochondrial structure, function, and metabolism and may partly contribute to many of the findings in patients. We found that antipsychotic CPZ was non-significantly associated with the cf-mtDNA measures. Cell-free mtDNA in the blood decreases progressively following certain antipsychotic treatments [20], suggesting that the elevated cf-mtDNA observed in medicated patients should not be primarily due to antipsychotic medications. The PRESS sequence with TE=30 ms is not optimized for lactate quantification, which ideally requires a longer TE (e.g., ∼144 ms) or spectral editing techniques to improve specificity, although our study aimed to evaluate a broad range of metabolites[90,91]. Future studies employing optimized sequences are needed to confirm these findings. The MRS was limited to one brain region - whether our findings can be applied to other brain regions is unclear. Our findings were based on cross-sectional studies, making it difficult to assess the causal pathway for the cf-mtDNA elevations in SSD. Our study also did not explore mtDNA mutation and other mitochondrial protein, gene expression, or morphological changes, leaving open the question on whether they are linked to the elevated cf-mtDNA in SSD. Also, the potential role of EV-derived mtDNA from brain cell types other than neurons, such as astrocytes and microglia, cannot be ruled out. MtDNA and nuclear DNA targets were not amplified in a multiplex qPCR reaction, which may introduce inter-assay variability and affect the accuracy of mtDNA quantification. To mitigate this, we normalized mtDNA levels to the average of three independent nuclear reference genes to enhance measurement robustness [66]. While our EV isolation strategy followed a validated protocol[70] and included immunoaffinity enrichment for neuronal L1CAM+ vesicles, we did not perform direct size characterization of the vesicles using techniques such as nanoparticle tracking analysis or electron microscopy. Future studies will include such biophysical assessments to further confirm EV purity and size distribution. Finally, time-of-day effects and acute lifestyle variables (e.g., recent exercise or dietary intake) that may influence cf-mtDNA levels were not systematically controlled, which should be considered in future studies.

In conclusion, peripherally obtained blood and particularly neuronal EV-based cf-mtDNA may provide a potential biomarker for indexing brain mitochondrial dysfunctions in SSD. Our initial exploration of how this putative biomarker is linked to the etiopathophysiology of SSD is encouraging as demonstrated by the significant association with childhood trauma in the patients. Cell-free mtDNA in the blood and in neuronal EVs may provide an intriguing window to study several aspects of the mitochondrial hypothesis such as the potential role of mitochondrial dysfunction in the stress-related psychosis process, the brain and body accelerated aging, and the high medical comorbidity issues in people with SSD.

## Data Availability Statement

The datasets generated and analyzed here are currently not publicly available, but we will strive to honor data request on case-by-case basis.

## Author Contributions

AA, AT, AKP and LEH wrote the paper, LEH and AKP obtained funding, AA, AT, BP, YM, JJC, SI, BA, PK contributed to data collection, processing and/or analysis, all contributed to manuscript editing, critical revision, and approved the final version of the manuscript.

## References

1. Whatley SA, Curti D, Marchbanks RM. Mitochondrial involvement in schizophrenia and other functional psychoses. Neurochem Res. 1996;21:995–1004.

2. Srivastava R, Faust T, Ramos A, Ishizuka K, Sawa A. Dynamic Changes of the Mitochondria in Psychiatric Illnesses: New Mechanistic Insights From Human Neuronal Models. Biol Psychiatry. 2018;83:751–760.

3. Whitehurst T, Howes O. The role of mitochondria in the pathophysiology of schizophrenia: a critical review of the evidence focusing on mitochondrial complex one. Neurosci Biobehav Rev. 2022;132:449–464.

4. Chouinard V-A, Du F, Chen X, Tusuzian E, Ren B, Anderson J, et al. Cognitive Impairment in Psychotic Disorders Is Associated with Brain Reductive Stress and Impaired Energy Metabolism as Measured by 31P Magnetic Resonance Spectroscopy. Schizophr Bull. 2025:sbaf003.

5. Du F, Cooper AJ, Thida T, Sehovic S, Lukas SE, Cohen BM, et al. In Vivo Evidence for Cerebral Bioenergetic Abnormalities in Schizophrenia Measured Using 31P Magnetization Transfer Spectroscopy. JAMA Psychiatry. 2014;71:19–27.

6. Roberts RC. Mitochondrial dysfunction in schizophrenia: With a focus on postmortem studies. Mitochondrion. 2021;56:91–101.

7. DiMauro S, Schon EA. Mitochondrial respiratory-chain diseases. New England Journal of Medicine. 2003;348:2656–2668.

8. Rollins B, Martin M V, Sequeira PA, Moon EA, Morgan LZ, Watson SJ, et al. Mitochondrial variants in schizophrenia, bipolar disorder, and major depressive disorder. PLoS One. 2009;4:e4913.

9. Bulduk BK, Tortajada J, Valiente-Pallejà A, Callado LF, Torrell H, Vilella E, et al. High number of mitochondrial DNA alterations in postmortem brain tissue of patients with schizophrenia compared to healthy controls. Psychiatry Res. 2024;337:115928.

10. Kumar P, Efstathopoulos P, Millischer V, Olsson E, Wei Y Bin, Brüstle O, et al. Mitochondrial DNA copy number is associated with psychosis severity and anti-psychotic treatment. Sci Rep. 2018;8:12743.

11. Verge B, Alonso Y, Valero J, Miralles C, Vilella E, Martorell L. Mitochondrial DNA (mtDNA) and schizophrenia. European Psychiatry. 2011;26:45–56.

12. Robicsek O, Karry R, Petit I, Salman-Kesner N, Müller FJ, Klein E, et al. Abnormal neuronal differentiation and mitochondrial dysfunction in hair follicle-derived induced pluripotent stem cells of schizophrenia patients. Mol Psychiatry. 2013;18:1067–1076.

13. Bertholet AM, Delerue T, Millet AM, Moulis MF, David C, Daloyau M, et al. Mitochondrial fusion/fission dynamics in neurodegeneration and neuronal plasticity. Neurobiol Dis. 2016;90:3– 19.

14. Shi R, Guberman M, Kirshenbaum LA. Mitochondrial quality control: the role of mitophagy in aging. Trends Cardiovasc Med. 2018;28:246–260.

15. Stephens OR, Grant D, Frimel M, Wanner N, Yin M, Willard B, et al. Characterization and origins of cell-free mitochondria in healthy murine and human blood. Mitochondrion. 2020;54:102–112.

16. Stier A. Human blood contains circulating cell-free mitochondria, but are they really functional? American Journal of Physiology-Endocrinology and Metabolism. 2021;320:E859–E863.

17. Pérez-Treviño P, Velásquez M, García N. Mechanisms of mitochondrial DNA escape and its relationship with different metabolic diseases. Biochimica et Biophysica Acta (BBA) - Molecular Basis of Disease. 2020;1866:165761.

18. García-De La Cruz DD, Juárez-Rojop IE, Tovilla-Zárate CA, Martínez-Magaña JJ, Genis-Mendoza AD, Nicolini H, et al. Association between mitochondrial DNA and cognitive impairment in schizophrenia: Study protocol for a mexican population. Neuropsychiatr Dis Treat. 2019;15:1717– 1722.

19. Li S, Jiang J, Zhu W, Wang D, Dong C, Bu Y, et al. Increased cell-free DNA is associated with oxidative damage in patients with schizophrenia. J Psychiatr Res. 2024;175:20–28.

20. Ouyang H, Huang M, Xu Y, Yao Q, Wu X, Zhou D. Reduced Cell-Free Mitochondrial DNA Levels Were Induced by Antipsychotics Treatment in First-Episode Patients With Schizophrenia. Front Psychiatry. 2021;12.

21. Yáñez-Mó M, Siljander PR-M, Andreu Z, Bedina Zavec A, Borràs FE, Buzas EI, et al. Biological properties of extracellular vesicles and their physiological functions. J Extracell Vesicles. 2015;4:27066.

22. Xue T, Liu W, Wang L, Shi Y, Hu Y, Yang J, et al. Extracellular vesicle biomarkers for complement dysfunction in schizophrenia. Brain. 2024;147:1075–1086.

23. Melentijevic I, Toth ML, Arnold ML, Guasp RJ, Harinath G, Nguyen KC, et al. C. elegans neurons jettison protein aggregates and mitochondria under neurotoxic stress. Nature. 2017;542:367– 371.

24. Goetzl EJ, Srihari VH, Guloksuz S, Ferrara M, Tek C, Heninger GR. Neural cell-derived plasma exosome protein abnormalities implicate mitochondrial impairment in first episodes of psychosis. The FASEB Journal. 2021;35:e21339.

25. Benarroch E. What Is the Role of Lactate in Brain Metabolism, Plasticity, and Neurodegeneration? Neurology. 2024;102:e209378.

26. Karagiannis A, Gallopin T, Lacroix A, Plaisier F, Piquet J, Geoffroy H, et al. Lactate is an energy substrate for rodent cortical neurons and enhances their firing activity. 2021. 2021. 10.7554/eLife.

27. Chiappelli J, Savransky A, Ma Y, Gao S, Kvarta MD, Kochunov P, et al. Impact of lifetime stressor exposure on neuroenergetics in schizophrenia spectrum disorders. Schizophr Res. 2024;269:58– 63.

28. Dogan AE, Yuksel C, Du F, Chouinard VA, Öngür D. Brain lactate and pH in schizophrenia and bipolar disorder: A systematic review of findings from magnetic resonance studies. Neuropsychopharmacology. 2018;43:1681–1690.

29. Rowland LM, Pradhan S, Korenic S, Wijtenburg SA, Hong LE, Edden RA, et al. Elevated brain lactate in schizophrenia: A 7T magnetic resonance spectroscopy study. Transl Psychiatry. 2016;6.

30. Liu S, Zhang L, Fan X, Wang G, Liu Q, Yang Y, et al. Lactate levels in the brain and blood of schizophrenia patients: A systematic review and meta-analysis. Schizophr Res. 2024;264:29–38.

31. Sullivan CR, Mielnik CA, Funk A, O’Donovan SM, Bentea E, Pletnikov M, et al. Measurement of lactate levels in postmortem brain, iPSCs, and animal models of schizophrenia. Sci Rep. 2019;9.

32. Nugent KL, Chiappelli J, Rowland LM, Daughters SB, Hong LE. Distress intolerance and clinical functioning in persons with schizophrenia. Psychiatry Res. 2014;220:31–36.

33. Da Silva T, Wu A, Laksono I, Prce I, Maheandiran M, Kiang M, et al. Mitochondrial function in individuals at clinical high risk for psychosis. Sci Rep. 2018;8.

34. Roberts RC, Barksdale KA, Roche JK, Lahti AC. Decreased synaptic and mitochondrial density in the postmortem anterior cingulate cortex in schizophrenia. Schizophr Res. 2015;168:543–553.

35. Akter M, Hasan M, Ramkrishnan AS, Iqbal Z, Zheng X, Fu Z, et al. Astrocyte and L-lactate in the anterior cingulate cortex modulate schema memory and neuronal mitochondrial biogenesis. Elife. 2023;12.

36. Zemirli N, Morel E, Molino D. Mitochondrial dynamics in basal and stressful conditions. Int J Mol Sci. 2018;19:564.

37. Picard M, McEwen BS. Psychological stress and mitochondria: a conceptual framework. Biopsychosocial Science and Medicine. 2018;80:126–140.

38. Van Os J, Selten J-P. Prenatal exposure to maternal stress and subsequent schizophrenia: the May 1940 invasion of the Netherlands. The British Journal of Psychiatry. 1998;172:324–326.

39. Khashan AS, Abel KM, McNamee R, Pedersen MG, Webb RT, Baker PN, et al. Higher risk of offspring schizophrenia following antenatal maternal exposure to severe adverse life events. Arch Gen Psychiatry. 2008;65:146–152.

40. Varese F, Smeets F, Drukker M, Lieverse R, Lataster T, Viechtbauer W, et al. Childhood adversities increase the risk of psychosis: a meta-analysis of patient-control, prospective-and cross-sectional cohort studies. Schizophr Bull. 2012;38:661–671.

41. Docherty NM, St-Hilaire A, Aakre JM, Seghers JP. Life events and high-trait reactivity together predict psychotic symptom increases in schizophrenia. Schizophr Bull. 2009;35:638–645.

42. Beards S, Gayer-Anderson C, Borges S, Dewey ME, Fisher HL, Morgan C. Life events and psychosis: a review and meta-analysis. Schizophr Bull. 2013;39:740–747.

43. Tripathi A, Bartosh A, Whitehead C, Pillai A. Activation of cell-free mtDNA-TLR9 signaling mediates chronic stress-induced social behavior deficits. Mol Psychiatry. 2023;28:3806–3815.

44. Krolow R, Noschang C, Weis SN, Pettenuzzo LF, Huffell AP, Arcego DM, et al. Isolation stress during the prepubertal period in rats induces long-lasting neurochemical changes in the prefrontal cortex. Neurochem Res. 2012;37:1063–1073.

45. Ridout KK, Coe JL, Parade SH, Marsit CJ, Kao H-T, Porton B, et al. Molecular markers of neuroendocrine function and mitochondrial biogenesis associated with early life stress. Psychoneuroendocrinology. 2020;116:104632.

46. Ruigrok SR, Yim K, Emmerzaal TL, Geenen B, Stöberl N, den Blaauwen JL, et al. Effects of early-life stress on peripheral and central mitochondria in male mice across ages. Psychoneuroendocrinology. 2021;132:105346.

47. Woods SW. Chlorpromazine equivalent doses for the newer atypical antipsychotics. Journal of Clinical Psychiatry. 2003;64:663–667.

48. Overall JE, Gorham DR. The brief psychiatric rating scale. Psychol Rep. 1962;10:799–812.

49. Kirkpatrick B, Strauss GP, Nguyen L, Fischer BA, Daniel DG, Cienfuegos A, et al. The brief negative symptom scale: psychometric properties. Schizophr Bull. 2011;37:300–305.

50. Bernstein DP, Stein JA, Newcomb MD, Walker E, Pogge D, Ahluvalia T, et al. Development and validation of a brief screening version of the Childhood Trauma Questionnaire. Child Abuse Negl. 2003;27:169–190.

51. Cohen S, Kamarck T, Mermelstein R. A global measure of perceived stress. J Health Soc Behav. 1983:385–396.

52. Mullins PG, McGonigle DJ, O’Gorman RL, Puts NAJ, Vidyasagar R, Evans CJ, et al. Current practice in the use of MEGA-PRESS spectroscopy for the detection of GABA. Neuroimage. 2014;86:43–52.

53. Yoon SH, Park CM, Lee CH, Song I-C, Lee HJ, Goo JM. Feasibility of In vivo Proton Magnetic Resonance Spectroscopy for Lung Cancer. Journal of the Korean Society of Magnetic Resonance in Medicine. 2012;16:40–46.

54. Rowland LM, Pradhan S, Korenic S, Wijtenburg SA, Hong LE, Edden RA, et al. Elevated brain lactate in schizophrenia: a 7 T magnetic resonance spectroscopy study. Transl Psychiatry. 2016;6:e967–e967.

55. Erbay MF, Zayman EP, Erbay LG, Ünal S. Evaluation of transcranial magnetic stimulation efficiency in major depressive disorder patients: A magnetic resonance spectroscopy study. Psychiatry Investig. 2019;16:745–750.

56. Soeiro-De-Souza MG, Pastorello BF, Da Costa Leite C, Henning A, Moreno RA, Otaduy MCG. Dorsal anterior cingulate lactate and glutathione levels in euthymic bipolar i disorder: 1H-MRS study. International Journal of Neuropsychopharmacology. 2016;19:1–8.

57. Batistuzzo MC, Sottili BA, Shavitt RG, Lopes AC, Cappi C, de Mathis MA, et al. Lower Ventromedial Prefrontal Cortex Glutamate Levels in Patients With Obsessive–Compulsive Disorder. Front Psychiatry. 2021;12.

58. Koush Y, de Graaf RA, Kupers R, Dricot L, Ptito M, Behar KL, et al. Metabolic underpinnings of activated and deactivated cortical areas in human brain. Journal of Cerebral Blood Flow and Metabolism. 2021;41:986–1000.

59. Xu J, Dydak U, Harezlak J, Nixon J, Dzemidzic M, Gunn AD, et al. Neurochemical Abnormalities in Unmedicated Bipolar Depression and Mania: A 2D 1H MRS Investigation. Psychiatry Res. 2013;213:235.

60. Quadrelli S, Mountford C, Ramadan S. Hitchhiker’S Guide to Voxel Segmentation for Partial Volume Correction of in Vivo Magnetic Resonance Spectroscopy. Magn Reson Insights. 2016;9:MRI.S32903.

61. Near J, Harris AD, Juchem C, Kreis R, Marjańska M, Öz G, et al. Preprocessing, analysis and quantification in single-voxel magnetic resonance spectroscopy: experts’ consensus recommendations. NMR Biomed. 2021;34.

62. Regenold WT, Phatak P, Marano CM, Sassan A, Conley RR, Kling MA. Elevated Cerebrospinal Fluid Lactate Concentrations in Patients with Bipolar Disorder and Schizophrenia: Implications for the Mitochondrial Dysfunction Hypothesis. Biol Psychiatry. 2008;65:489.

63. Provencher SW. Automatic quantitation of localized in vivo 1H spectra with LCModel. NMR Biomed. 2001;14:260–264.

64. Zöllner HJ, Davies-Jenkins CW, Murali-Manohar S, Gong T, Hui SCN, Song Y, et al. Feasibility and implications of using subject-specific macromolecular spectra to model short-TE MRS data. NMR Biomed. 2022;36:e4854.

65. Trumpff C, Michelson J, Lagranha CJ, Taleon V, Karan KR, Sturm G, et al. Stress and circulating cell-free mitochondrial DNA: A systematic review of human studies, physiological considerations, and technical recommendations. Mitochondrion. 2021;59:225–245.

66. Maggo S, North LY, Ozuna A, Ostrow D, Grajeda YR, Hakimjavadi H, et al. A method for measuring mitochondrial DNA copy number in pediatric populations. Front Pediatr. 2024;12.

67. Livak KJ, Schmittgen TD. Analysis of relative gene expression data using real-time quantitative PCR and the 2-ΔΔCT method. Methods. 2001;25:402–408.

68. Zhang L, Deng S, Zhao S, Ai Y, Zhang L, Pan P, et al. Intra-peritoneal administration of mitochondrial DNA provokes acute lung injury and systemic inflammation via toll-like receptor 9. Int J Mol Sci. 2016;17.

69. Wiersma M, van Marion DMS, Bouman EJ, Li J, Zhang D, Ramos KS, et al. Cell-free circulating mitochondrial dna: A potential blood-based marker for atrial fibrillation. Cells. 2020;9.

70. Mustapic M, Eitan E, Werner JK, Berkowitz ST, Lazaropoulos MP, Tran J, et al. Plasma extracellular vesicles enriched for neuronal origin: A potential window into brain pathologic processes. Front Neurosci. 2017;11.

71. Neustadt J, Pieczenik SR. Medication-induced mitochondrial damage and disease. Mol Nutr Food Res. 2008;52:780–788.

72. Chan ST, McCarthy MJ, Vawter MP. Psychiatric drugs impact mitochondrial function in brain and other tissues. Schizophr Res. 2019;217:136.

73. Cakir Y, Yang Z, Knight CA, Pompilius M, Westbrook D, Bailey SM, et al. Effect of alcohol and tobacco smoke on mtDNA damage and atherogenesis. Free Radic Biol Med. 2007;43:1279–1288.

74. Yang Z, Harrison CM, Chuang GC, Ballinger SW. The role of tobacco smoke induced mitochondrial damage in vascular dysfunction and atherosclerosis. Mutation Research/Fundamental and Molecular Mechanisms of Mutagenesis. 2007;621:61–74.

75. Norman M, Ter-Ovanesyan D, Trieu W, Lazarovits R, Kowal EJK, Lee JH, et al. L1CAM is not associated with extracellular vesicles in human cerebrospinal fluid or plasma. Nat Methods. 2021;18:631–634.

76. Nogueras-Ortiz CJ, Eren E, Yao P, Calzada E, Dunn C, Volpert O, et al. Single-extracellular vesicle (EV) analyses validate the use of L1 cell adhesion molecule (L1CAM) as a reliable biomarker of neuron-derived EVs. J Extracell Vesicles. 2024;13:e12459.

77. Sprenger H-G, Langer T. The good and the bad of mitochondrial breakups. Trends Cell Biol. 2019;29:888–900.

78. Santos C, Martinez M, Lima M, Hao Y-J, Simoes N, Montiel R. Mitochondrial DNA mutations in cancer: a review. Curr Top Med Chem. 2008;8:1351–1366.

79. Smith ALM, Whitehall JC, Greaves LC. Mitochondrial DNA mutations in ageing and cancer. Mol Oncol. 2022;16:3276–3294.

80. Khotina VA, Vinokurov AY, Sinyov V V, Zhuravlev AD, Popov DY, Sukhorukov VN, et al. Mitochondrial dysfunction associated with mtDNA mutation: mitochondrial genome editing in atherosclerosis research. Curr Med Chem. 2024. 2024.

81. Risi B, Imarisio A, Cuconato G, Padovani A, Valente EM, Filosto M. Mitochondrial DNA (mtDNA) as fluid biomarker in neurodegenerative disorders: A systematic review. Eur J Neurol. 2025;32.

82. Fuke S, Kametani M, Kato T. Quantitative analysis of the 4977-bp common deletion of mitochondrial DNA in postmortem frontal cortex from patients with bipolar disorder and schizophrenia. Neurosci Lett. 2008;439:173–177.

83. Mamdani F, Rollins B, Morgan L, Sequeira PA, Vawter MP. The somatic common deletion in mitochondrial DNA is decreased in schizophrenia. Schizophr Res. 2014;159:370–375.

84. Win PW, Singh SM, Castellani CA. Mitochondrial DNA Copy Number and Heteroplasmy in Monozygotic Twins Discordant for Schizophrenia. Twin Research and Human Genetics. 2023;26:280–289.

85. San-Millan I, Sparagna GC, Chapman HL, Warkins VL, Chatfield KC, Shuff SR, et al. Chronic Lactate Exposure Decreases Mitochondrial Function by Inhibition of Fatty Acid Uptake and Cardiolipin Alterations in Neonatal Rat Cardiomyocytes. Front Nutr. 2022;9.

86. Glancy B, Kane DA, Kavazis AN, Goodwin ML, Willis WT, Gladden LB. Mitochondrial lactate metabolism: history and implications for exercise and disease. Journal of Physiology. 2021;599:863–888.

87. Takahashi K, Tamura Y, Kitaoka Y, Matsunaga Y, Hatta H. Effects of Lactate Administration on Mitochondrial Respiratory Function in Mouse Skeletal Muscle. Front Physiol. 2022;13.

88. Brooks GA, Dubouchaud H, Brown M, Sicurello JP, Butz CE. Role of mitochondrial lactate dehydrogenase and lactate oxidation in the intracellular lactate shuttle. vol. 96. 1999.

89. Kochunov P, Ma Y, Hatch KS, Gao S, Acheson A, Jahanshad N, et al. Ancestral, Pregnancy, and Negative Early-Life Risks Shape Children’s Brain (Dis)similarity to Schizophrenia. Biol Psychiatry. 2023;94:332–340.

90. Robison RK, Haynes JR, Ganji SK, Nockowski CP, Kovacs Z, Pham W, et al. J-Difference editing (MEGA) of lactate in the human brain at 3T. Magn Reson Med. 2023;90:852–862.

91. Maier S, Nickel K, Lange T, Oeltzschner G, Dacko M, Endres D, et al. Increased cerebral lactate levels in adults with autism spectrum disorders compared to non-autistic controls: a magnetic resonance spectroscopy study. Mol Autism. 2023;14.

