## Supplementary Information for "Blood and Neuronal Extracellular Vesicle Mitochondrial Disruptions in Schizophrenia"

### SI1. MtDNA genotyping.

Primers were synthesized by Integrated DNA Technologies. MasterCycler (Eppendorf, Westburg, NY, USA) was used to perform qRT-PCR using iTaqTM Universal SYBR® Green Supermix (Bio-Rad, CA, USA). Mitochondrial DNA levels were adjusted for nuclear DNA (nDNA) levels using the average values of three nuclear genes; 18 S rRNA (forward primer-5' AGAAACGGCTACCACATCCA 3' and reverse primer-5' CCCTCCAATGGATCCTCGTT 3'), B2M (forward primer-5' CACTGAAAAAGATGAGTATGCC 3' and reverse primer-5' AACATTCCCTGACAATCCC 3'), and β-actin (forward primer-5' GGACTTCGAGCAAGAGATGG 3' and reverse primer-5' AGCACTGTGTTGGCGTACAG 3').

### SI2. Total EV isolation.

EVs were isolated from 0.5ml of human serum following a detailed protocol where 485µl of Dulbecco's calcium- and magnesium-free salt solution was added, supplemented with protease and phosphatase inhibitor cocktail (#78440, Thermo Fisher Scientific) at three times the recommended concentrations. Samples were mixed, left at room temperature for 5 minutes, and then centrifuged at 4,000×g for 20 minutes at 4°C. The supernatants were transferred to fresh tubes, gently mixed by inversion, and 252µl of ExoQuick™ exosome precipitation solution (System Biosciences, Mountain View, CA) was added. After a 60-minute incubation at 4°C, the samples were centrifuged at 1,500×g for 20 minutes at 4°C to pellet total EVs. The supernatants were discarded, and the pellets were re-suspended in 0.5ml of ultrapure distilled water (Invitrogen-Thermo Fisher Scientific, Rockford, IL) containing protease and phosphatase inhibitors at three times the recommended concentrations.

**SI3. Neuronal EV isolation and validation.**

To enrich for neuronal EVs from the total EV, the suspensions were incubated at 4°C for 1 hour with 4 µg of biotinylated anti-human L1 cell adhesion molecule (L1CAM) antibody (clone 5G3, eBioscience, San Diego, CA) in 50µl of 3% BSA solution (Blocker BSA 10% solution in PBS, Thermo Scientific, Rockford, IL) per tube with continuous mixing. Streptavidin-agarose Ultralink resin (15 µl; Thermo Scientific, Rockford, IL) in 40µl of 3% BSA solution was then added, and the mixture was incubated for 30 minutes at 4°C with continuous bound EVs. Pellets were re-suspended in 200 µl of 0.1 M glycine-HCl solution, mixed for 10 seconds, and centrifuged at 4,500×g for 10 minutes at 4°C to detach the neuronal EVs from the bead-antibody complex. The supernatants were transferred to clean tubes containing 25µl of 10% BSA and 15µl of 1M Tris-HCl and mixed thoroughly.

To validate the efficiency of the L1CAM pulldown, we analyzed the expression of L1CAM in neuronal EVs comparing with total EVs. To evaluate the neuronal enrichment of blood-derived neuronal EVs, we examined the protein expression of the neuronal marker microtubule-associated protein-2 (MAP2), which has two isoforms: high molecular weight

(MAP2 A/B) and low molecular weight (MAP2 C/D) by immunoblot analysis. We also analyzed the canonical EV marker CD9 [1,2] in these samples. Human brain lysate and mouse spleen lysate were used as controls [3]. Briefly, to lyse the EVs, 260µl of mammalian protein extraction reagent (M-PER; Thermo Scientific, Rockford, IL, USA) supplemented with protease and phosphatase inhibitors at three times the recommended concentrations was added to each tube. The samples underwent two freeze-thaw cycles to ensure complete lysis. The final suspensions containing EV proteins were stored at −80°C.

The protein samples were then separated via 4–15% SDS-PAGE gel electrophoresis and then transferred to PVDF membranes (#IPVH00010, Millipore). After the transfer, PVDF membranes were blocked with 5% nonfat dry milk in 1×TBST buffer [1.21g Tris (BP152-5, Fisher Scientific), 8.77g NaCl (#BP358-212, Fisher Scientific), 500 µL Tween-20 (#BP337-500, Fisher Scientific), pH 7.6 for 1 L]. Membranes were incubated with monoclonal antibodies against L1 cell adhesion molecule [L1CAM (C-2), sc-514360], MAP-2(C-2, sc-390543), and CD9 (H-110, sc-9148) at 4 °C overnight. After washing, membranes were incubated with the secondary antibodies conjugated with horseradish peroxidase (HRP) for 1 h followed by washing and detection using Super Signal West Pico Chemiluminescent Substrate (#34078, Thermo Fisher Scientific).

**SI4. Additional Results and Discussion on Lactate+**

The lactate spectra measured at 3T scanner using TE at 30 ms overlaps with lipid resonances, particularly in the ~1.3 ppm region [4]. The use of 30 ms TE does not allow adequate separation between lactate and these other metabolites, and thus the lactate measured here was labeled lactate+ to indicate that it may be contaminated with other metabolites. However, giving the interesting correlation findings between lactate+ and cf-mtDNA measures, we explored the level of lipid contamination effects by comparing lactate+ and lipid separately in their associations with cf-mtDNA markers (**SI Table 1**).

*SI Table 1. Correlation between anterior cingulate cortex (ACC) lactate+ and lipid metabolite levels and cf-mtDNA markers from neuronally-enriched extracellular vesicles (NEV) and blood in schizophrenia spectrum disorders (SSD) and healthy controls (HC).*

| | **SSD** | | **HC** | |
|---|---|---|---|---|
| | r | p | r | p |
| **NEV cf-mtDNA** | | | | |
| **Lacate+** | 0.53 | 0.03 | -0.28 | 0.17 |
| **Lip13a** | -0.04 | 0.90 | 0.11 | 0.74 |
| **Lip09** | 0.42 | 0.12 | 0.17 | 0.61 |
| **Lip13a.Lip13b** | -0.04 | 0.90 | 0.11 | 0.74 |
| **Blood cf-mtDNA** | | | | |
| **Lacate+** | 0.53 | 0.008 | -0.09 | 0.06 |
| **Lip13a** | -0.43 | 0.08 | 0.51 | 0.06 |
| **Lip09** | -0.17 | 0.49 | 0.52 | 0.06 |
| **Lip13a.Lip13b** | -0.43 | 0.08 | 0.51 | 0.06 |

The data showed that cf-mtDNA markers in both nEV and blood were significantly correlated with lactate+ and only in patients, but not for any of the lipid spectra in either patients or controls, suggesting that the lactate+ association findings are more likely lactate-related.

The MRS acquisition using short TE of 30 ms provides high signal-to-noise ratio (SNR) and minimal T2 signal decay, enabling reliable detection of several metabolites including J-coupled metabolites such as lactate [5,6]. Importantly, at this short TE the doublet characteristic of lactate at 1.3 ppm remains distinguishable, especially in phased spectra, allowing it to be partially resolved from broader lipid signals[7]. Unlike longer TEs (e.g., 144 ms), which may suppress lipid signal but also reduce lactate visibility, TE = 30 ms preserves signal amplitude and metabolite integrity [7,8]. We also used careful modeling of overlapping peaks using the validated spectral fitting too LCModel, which incorporates prior knowledge to distinguish lactate from lipid resonances, particularly when SNR is high. This also helps partially address the concern of misattributing lipid signal to lactate.

### References


1. Mustapic M, Eitan E, Werner JK, Berkowitz ST, Lazaropoulos MP, Tran J, et al. Plasma extracellular vesicles enriched for neuronal origin: A potential window into brain pathologic processes. Front Neurosci. 2017;11.
2. Helwa I, Cai J, Drewry MD, Zimmerman A, Dinkins MB, Khaled ML, et al. A comparative study of serum exosome isolation using differential ultracentrifugation and three commercial reagents. PLoS One. 2017;12.
3. Jalava NS, Lopez-Picon FR, Kukko-Lukjanov TK, Holopainen IE. Changes in microtubule-associated protein-2 (MAP2) expression during development and after status epilepticus in the immature rat hippocampus. International Journal of Developmental Neuroscience. 2007;25:121–131.
4. Kreis R. The trouble with quality filtering based on relative C ramér-R ao lower bounds. Magn Reson Med. 2016;75:15–18.
5. Öz G, Deelchand DK, Wijnen JP, Mlynárik V, Xin L, Mekle R, et al. Advanced single voxel 1H magnetic resonance spectroscopy techniques in humans: Experts' consensus recommendations. NMR Biomed. 2021;34.
6. Öz G, Tkáč I. Short-echo, single-shot, full-intensity proton magnetic resonance spectroscopy for neurochemical profiling at 4 T: Validation in the cerebellum and brainstem. Magn Reson Med. 2011;65:901–910.
7. Yamasaki F, Takaba J, Ohtaki M, Abe N, Kajiwara Y, Saito T, et al. Detection and differentiation of lactate and lipids by single-voxel proton MR spectroscopy. Neurosurg Rev. 2005;28:267–277.
8. Bogner W, Hangel G, Esmaeili M, Andronesi OC. 1D-spectral editing and 2D multispectral in vivo 1H-MRS and 1H-MRSI - Methods and applications. Anal Biochem. 2017;529:48–64.

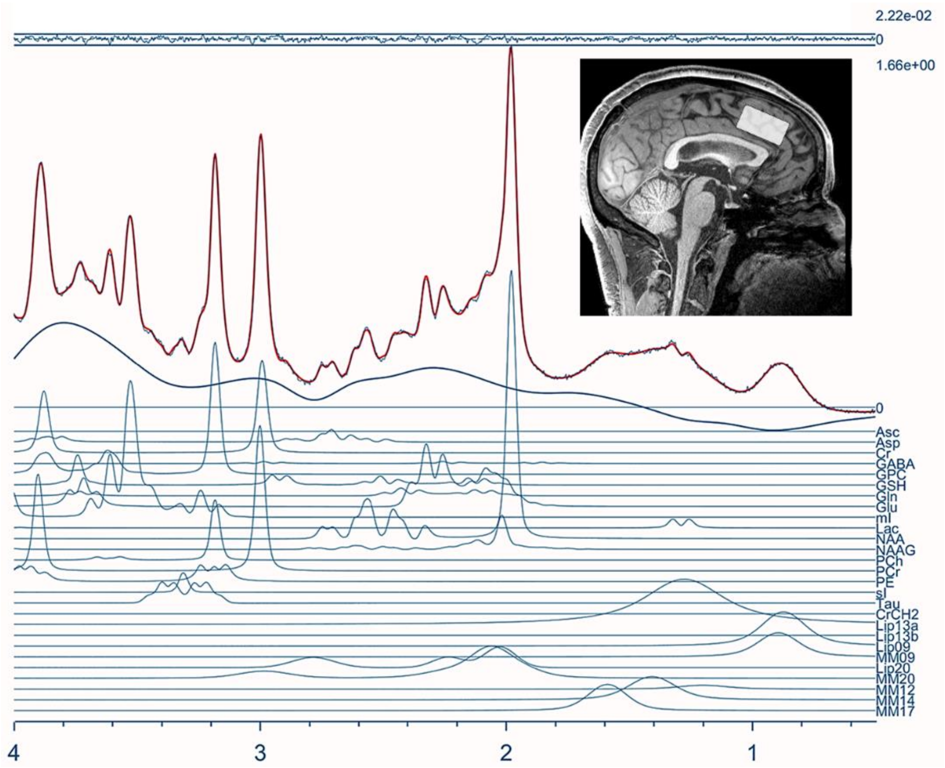


Supplementary Figure 1. Model Fit by using LC model basis set to identify the metabolite peaks.
